# The guardians of mitochondrial dynamics: a novel role for intermediate filament proteins

**DOI:** 10.1101/2024.07.19.604282

**Authors:** Irene MGM Hemel, Carlijn Steen, Simon LIJ Denil, Gökhan Ertaylan, Martina Kutmon, Michiel Adriaens, Mike Gerards

## Abstract

Mitochondria are dynamic organelles and the main source of cellular energy. Their dynamic nature is crucial to meet cellular requirements. However, the processes and proteins involved in mitochondrial dynamics are not fully understood. Using a computational protein-protein interaction approach, we identified ITPRIPL2, which caused mitochondrial elongation upon knockdown. ITPRIPL2 co-localizes with the intermediate filament protein vimentin and interacts with vimentin according to protein simulations. ITPRIPL2 knockdown alters vimentin processing, disrupts intermediate filaments and transcriptomics analysis revealed changes in vimentin-related pathways. Our data illustrates that ITPRIPL2 is essential for vimentin related intermediate filament structure. Interestingly, like ITPRIPL2 knockdown, vimentin knockdown results in mitochondrial elongation. Our data highlights ITPRIPL2 as a vimentin-associated protein and reveals a role for intermediate filaments in mitochondrial dynamics, improving our understanding of mitochondrial dynamics regulators. Moreover, our study demonstrates that protein-protein interaction analysis is a powerful approach for identifying novel mitochondrial dynamics proteins.

## 1 Introduction

Mitochondria are multifunctional, dynamic cellular organelles, crucial for a multitude of biochemical processes including cellular energy production and calcium homeostasis [1]. They display highly variable morphologies as the result of controlled cycles of fission and fusion, mitochondrial transport and mitophagy, collectively referred to as mitochondrial dynamics. Proper functioning and regulation of these processes is necessary to meet cellular energy requirements and to maintain a healthy mitochondrial pool. Mitochondrial fission results in the formation of small mitochondria suitable for transport and recycling of damaged mitochondria. The process of mitochondrial fission is driven by mitochondrial contact with the endoplasmatic reticulum (ER) and recruitment of dynamin-1-like protein (Drp1) [1, 2]. However, this process is more complex and requires many additional proteins, like mitochondrial fission factor (MFF) and mitochondrial dynamics protein MID51, as well as the polymerization of short actin filaments [2–4]. Transport of mitochondria is dependent on mitochondrial fission and ensures that energy requirements are met in all cellular compartments [1]. Movement of mitochondria to and from the cellular sections that require energy is necessary in all cell types and is especially important for mitochondrial placement in neurons [5]. Mitochondrial fusion, is the result of subsequent outer and inner membrane fusion induced by mitofusin 1/2 (Mfn1/2) as well as dynamin-like 120 kDa protein (OPA1) [2]. Mitochondrial fusion enables exchange of mitochondrial content and prevents degradation through mitophagy. PINK1/Parkin dependent mitophagy is crucial for mitochondrial quality control and removes damaged mitochondria [1]. Under physiological conditions, the balance between the different mitochondrial dynamics processes is tightly regulated and determines mitochondrial shape, size, localization and number [6]. This balance varies between cell and tissue types, with short spherical mitochondria in for instance hepatocytes and longer tubular mitochondria in fibroblasts, depending on subcellular energy demand [7, 8]. Changes in the balance can occur upon cellular stress, variability in nutrient availability or defects in fission or fusion processes which results in suboptimal mitochondrial morphology [2, 9]. Additionally, different organelles have an effect on the mitochondrial dynamics processes, including the ER, lysosomes and microtubules [4, 10, 11]. Changes in these organelles can alter mitochondrial morphology and impair optimal functioning.

Various disorders have been associated with mitochondrial dynamics, including Charcot-Marie-Tooth disease, which can be caused by defects in *MFN2*, dominant optic atrophy as the result of mutations in *OPA1* and peripheral neuropathy resulting from mutations in *DNM1L* [12, 13]. Furthermore, altered mitochondrial morphologies have been found in multiple neurodegenerative disorders, including Alzheimer’s and Huntington’s disease, where mitochondrial fragmentation is observed [14, 15].

While many different proteins involved in mitochondrial dynamics have been identified over the years, the complete processes and regulatory mechanisms are currently not fully understood. For example, while hypothesis have been proposed, it is still uncertain how mitochondrial pre-constriction occurs at mitochondrial-ER contact sites [4]. Although new factors that influence mitochondrial dynamics have been identified in several disorders [16], in other diseases, such as Alzheimer’s, it is unclear what the driving factor behind the altered mitochondrial morphology is. Gaining a better understanding of the genes and mechanisms behind mitochondrial dynamics will lead to increased insight into the different disorders, which is required for the development of treatments.

In this study, we aimed to identify proteins and processes involved in mitochondrial dynamics by using a computational approach. Computational methods allow the interpretation and analysis of large amounts of data as well as the combination of data from multiple resources and are increasingly used to improve the understanding of complex biological systems [17]. In this study, we applied a protein-protein interaction network analysis, which identified novel candidates that change mitochondrial morphology upon knockdown. One of these candidates, inositol 1,4,5-trisphosphate receptor interacting protein-like 2 (ITPRIPL2), is an intermediate filament associated protein and we show an essential role for the intermediate filaments in maintaining mitochondrial morphology.

## 2 Results

### 2.1 Protein-Protein interaction network analysis reveals novel candidates

For candidate identification, a protein-protein interaction network was created with eighteen known mitochondrial dynamics proteins (Table 1), which was subsequently expanded with a set of proteins most closely connected to the query list based on the total connectivity of a protein to the query set, compared to their overall connectivity in the database. Networks were created with varying combinations of confidence scores (Figure S1). High confidence scores resulted in networks with subclusters of proteins, mostly interacting with each other instead of the query proteins, while a low score resulted in inclusion of interactions with limited evidence potentially leading to the addition of more false positives. The most suitable network was selected with a medium confidence score of 0.5 and expansion with 100 proteins (Figure 1A), which resulted in a densely connected network. Leave-one-out validation showed a robust network in which 72% of the query proteins re-occur in the network upon expansion when left out of the initial query search (Figure 1B). Not re-occurring proteins are mostly present on the outside of the network and have either none or one interaction with other query proteins, whereas the re-occurring proteins generally have multiple (in)direct interactions with other query proteins. To verify that the created network contains more mitochondrial proteins than expected by chance, an overrepresentation analysis was performed. The network contained a significant overrepresentation of mitochondrial proteins (p<0.001), with 42% of expanded proteins annotated as mitochondrial (Figure 1C).

**Figure 1:**
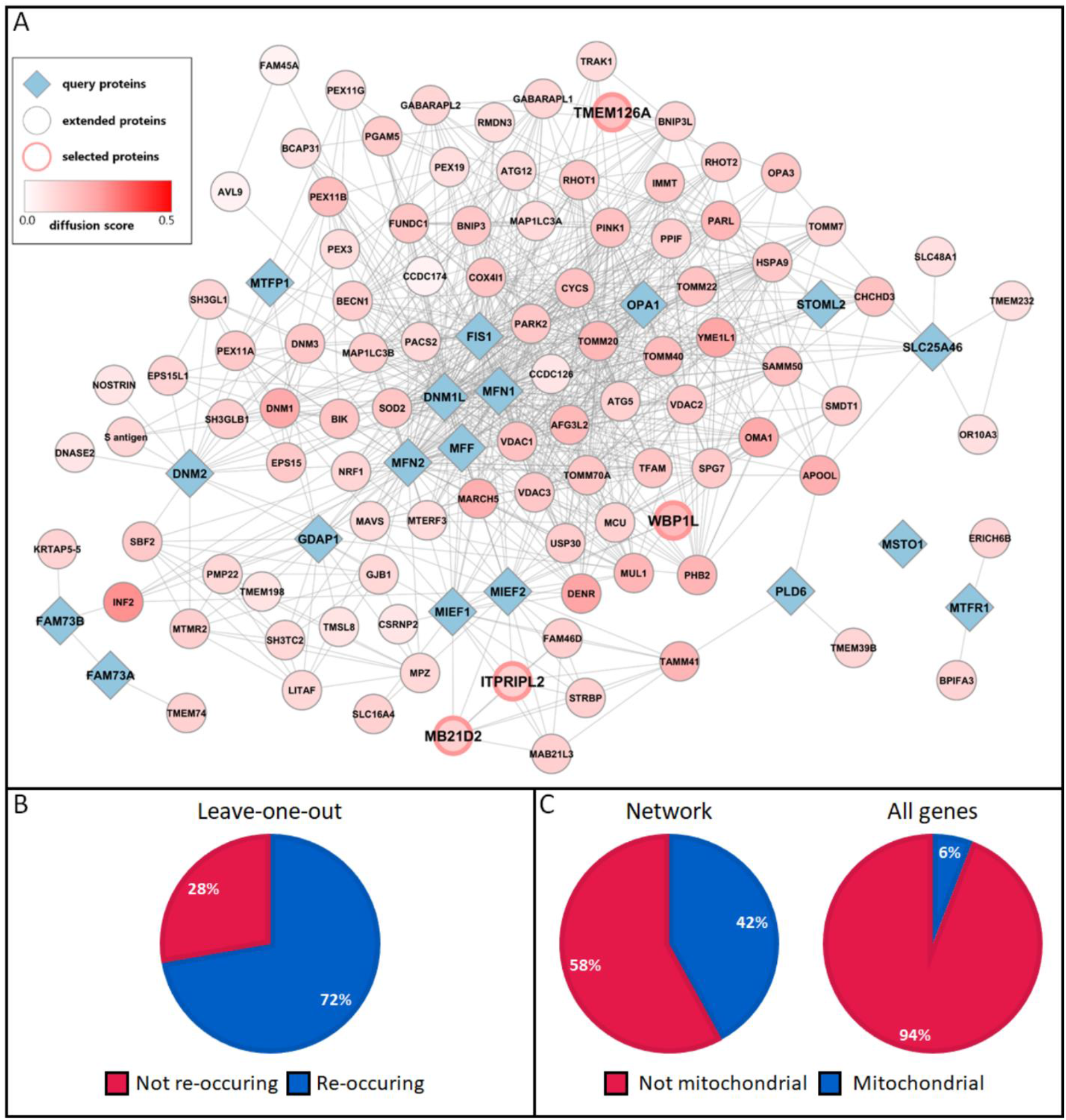
Protein-protein interaction network and validation. A) Protein-protein interaction network displaying the query proteins (diamond) and their known interactions with other proteins (ellipse). Heat propagation analysis is visualised on the node (colour gradient from white to red) where a higher score is indicative of proteins closer to the input proteins. Four proteins with no currently known function were selected for further analysis (red edged ellipses). B) Leave-one-out validation showed that 72% of the query proteins re-occur in the expanded network. C) Distribution of mitochondrial localisation for all expanded proteins. A significant overrepresentation of mitochondrial proteins was found in the network (p<0.001). The distribution of all genes in MitoCarta3.0 is shown for comparison.

**Table 1:** List of mitochondrial dynamics query proteins used for network generation.

| Gene | Protein | Uniprot identifier | Reference |
| --- | --- | --- | --- |
| *DNM1L* | Dynamin-1-like protein | O00429 | [18] |
| *DNM2* | Dynamin-2 | P50570 | [19] |
| *MFF* | Mitochondrial fission factor | Q9GZY8 | [20] |
| *MIEF2* | Mitochondrial dynamics protein MID49 | Q96C03 | [20] |
| *MIEF1* | Mitochondrial dynamics protein MID51 | Q9NQG6 | [20] |
| *FIS1* | Mitochondrial fission 1 protein | Q9Y3D6 | [20] |
| *GDAP1* | Ganglioside-induced differentiation-associated protein 1 | Q8TB36 | [21] |
| *SLC25A46* | Solute carrier family 25 member 46 | Q96AG3 | [22, 23] |
| *MTFP1* | Mitochondrial fission process protein 1 | Q9UDX5 | [24, 25] |
| *MTFR1* | Mitochondrial fission regulator 1 | Q15390 | [26] |
| *OPA1* | Dynamin-like 120 kDa protein | O60313 | [27, 28] |
| *MFN1* | Mitofusin-1 | Q8IWA4 | [29] |
| *MFN2* | Mitofusin-2 | O95140 | [29] |
| *MSTO1* | Protein misato homolog 1 | Q9BUK6 | [30, 31] |
| *MIGA1* | Mitoguardin 1 | Q8NAN2 | [32] |
| *MIGA2* | Mitoguardin 2 | Q7L4E1 | [32] |
| *PLD6* | Mitochondrial cardiolipin hydrolase | Q8N2A8 | [32, 33] |
| *STOML2* | Stomatin-like protein 2 | Q9UJZ1 | [34, 35] |

Heat propagation analysis was used as a measure of connectivity between query and candidate proteins. Within the twenty highest-ranking proteins, nine have a previously established association with mitochondrial dynamics, indicating that proteins with a link to mitochondrial dynamics can be predicted with this approach. In total, fifteen candidate proteins had a previous association with mitochondrial dynamics and were therefore excluded from further functional analysis (Table S1). For thirteen proteins, no function was reported at the time of candidate selection. Of these, the four highest-ranking candidates, expressed in fibroblasts, were selected for further functional validation; WBP1L, TMEM126A, MB21D2 and ITPRIPL2.

### 2.2 Knockdown of candidates alters mitochondrial morphology

A transient esiRNA mediated knockdown was created for the four selected candidates (Figure 2A). Knockdown efficiency was assessed and all candidate knockdown conditions showed a decrease in gene expression between 50 and 90% (Figure S2A). Upon knockdown, there was a significant increase in mean mitochondrial area for all four candidates, indicative of an altered balance between fission and fusion (Figure 2B). Additionally, mitochondrial branching was analysed as an indication of mitochondrial complexity. An increase in the number of branches per mitochondria, as well as an increase in the total branch length per mitochondrion was observed for WBP1L, TMEM126A, MB21D2 and ITPRIPL2, indicating an increased mitochondrial network complexity (Figure S2B/C). Furthermore, the mean form factor, a parameter that provides information on mitochondrial shape, showed significant differences between all knockdown conditions and control, again indicating an increase in mitochondrial network complexity (Figure S2D).

**Figure 2:**
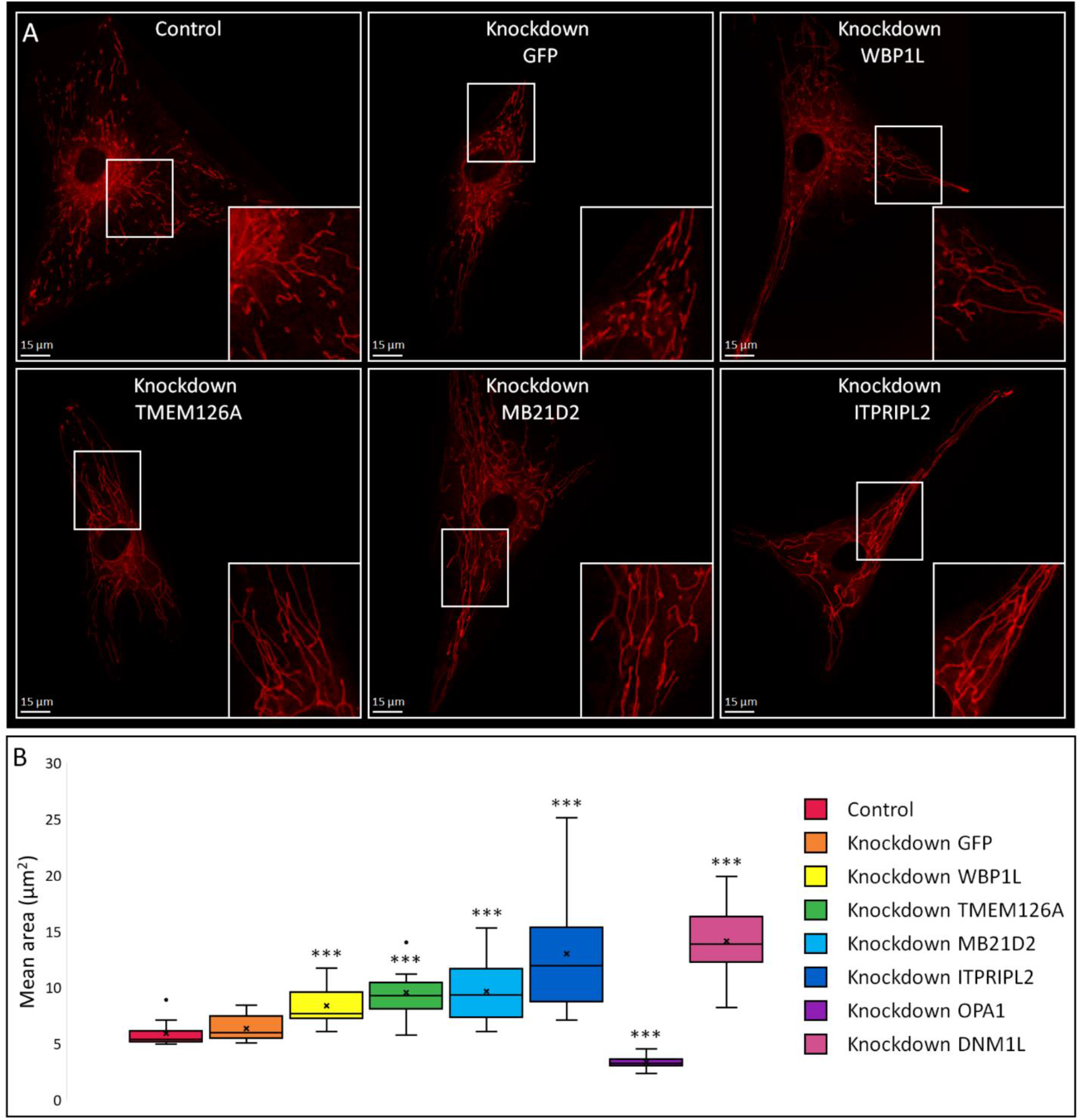
Assessment of mitochondrial morphology upon candidate knockdown. A) Examples of mitochondrial network morphology for control and knockdown conditions. B) Distribution of the mean mitochondrial area for control and knockdown conditions in human fibroblasts. A significant increase in mean area was observed for knockdown of WBP1L, TMEM126A, MB21D2 and ITPRIPL2. For positive controls OPA1 and DNM1L, a significant decrease and increase in mean area was observed, respectively, while mitochondrial morphology was unchanged for negative control GFP. Data information: Scale bars indicate 15 µm. Control N=18, Knockdown GFP, MB21D2 N=16, Knockdown WBP1L, TMEM126A, ITPRIPL2, OPA1 N=15, Knockdown DNM1L N=13. ***p < 0.001 (Mann Whitney U Test, significance level adjusted for multiple testing).

### 2.3 ITPRIPL2 is associated with the intermediate filaments

Since knockdown of ITPRIPL2 resulted in the most severe mitochondrial phenotype of the four candidates assessed, we decided to investigate the function of this protein in more detail. Subcellular localisation studies revealed that ITPRIPL2 co-localises with intermediate filament protein vimentin (Figure 3A), while no co-localisation with mitochondria was observed (Figure 3B). Interestingly, compression of the vimentin network occurs upon overexpression, indicating an effect of ITPRIPL2 presence on the structure of the intermediate filaments (Figure 3C). Furthermore, vimentin structure is disrupted upon ITPRIPL2 knockdown in fibroblasts, with fewer fibres present compared to non-transfected cells and clusters of vimentin appearing in the majority of ITPRIPL2 knockdown cells (Figure 3D). Although mRNA levels and western blot analysis did not show a change of total vimentin levels upon ITPRIPL2 knockdown, a shift in the ratio between the two vimentin forms was apparent with an increased abundance of the smaller form (Figure 4A/B). This implies that the disruption of the vimentin network structure caused by ITPRIPL2 knockdown is not the result of a decrease in vimentin levels, but by a shift in the type of vimentin present. Taken together, these results imply that ITPRIPL2 is involved in either vimentin processing or integration of vimentin proteins into intermediate filament fibres.

**Figure 3:**
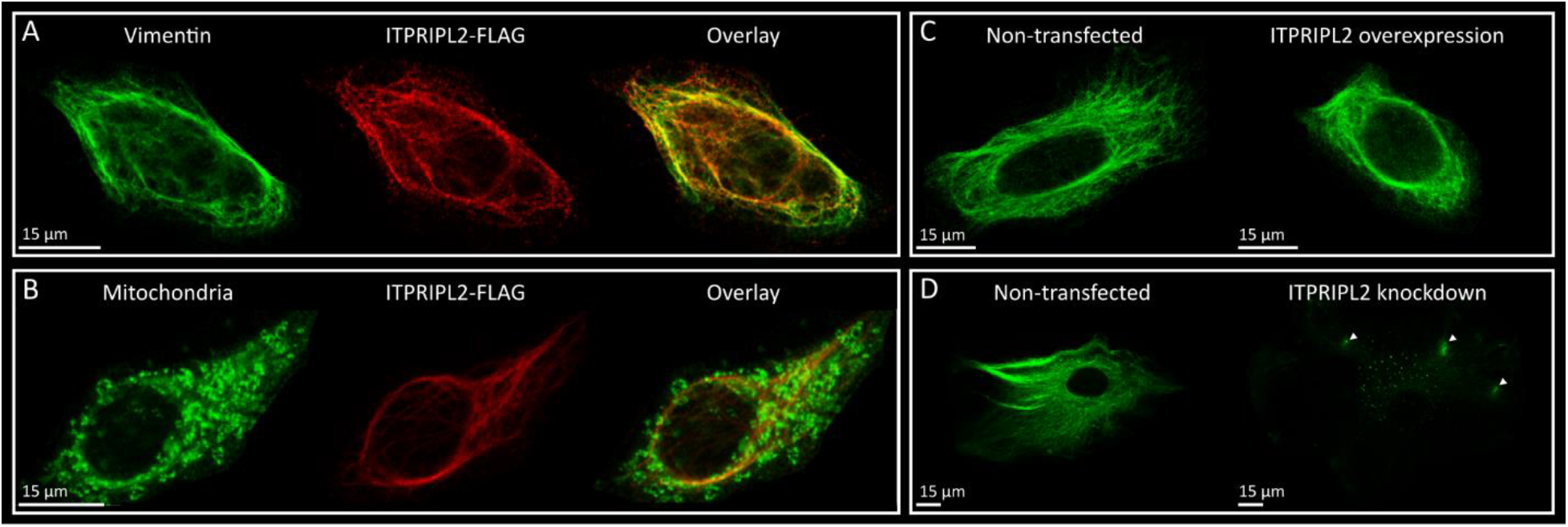
Co-localisation of ITPRIPL2. A) Immunostaining of ITPRIPL2 shows a partial co-localisation with vimentin, primarily surrounding the nucleus. B) ITPRIPL2 does not co-localise with mitochondria. C) Overexpression of ITPRIPL2 in HeLa cells results in condensation of the vimentin network around the nucleus compared to non-transfected cells. D) Knockdown of ITPRIPL2 in normal human dermal fibroblasts (nHDF) results in a decreased number of vimentin fibres and clusters of vimentin across the cell (white arrows). Data information: Scale bars indicate 15 µm.

**Figure 4:**
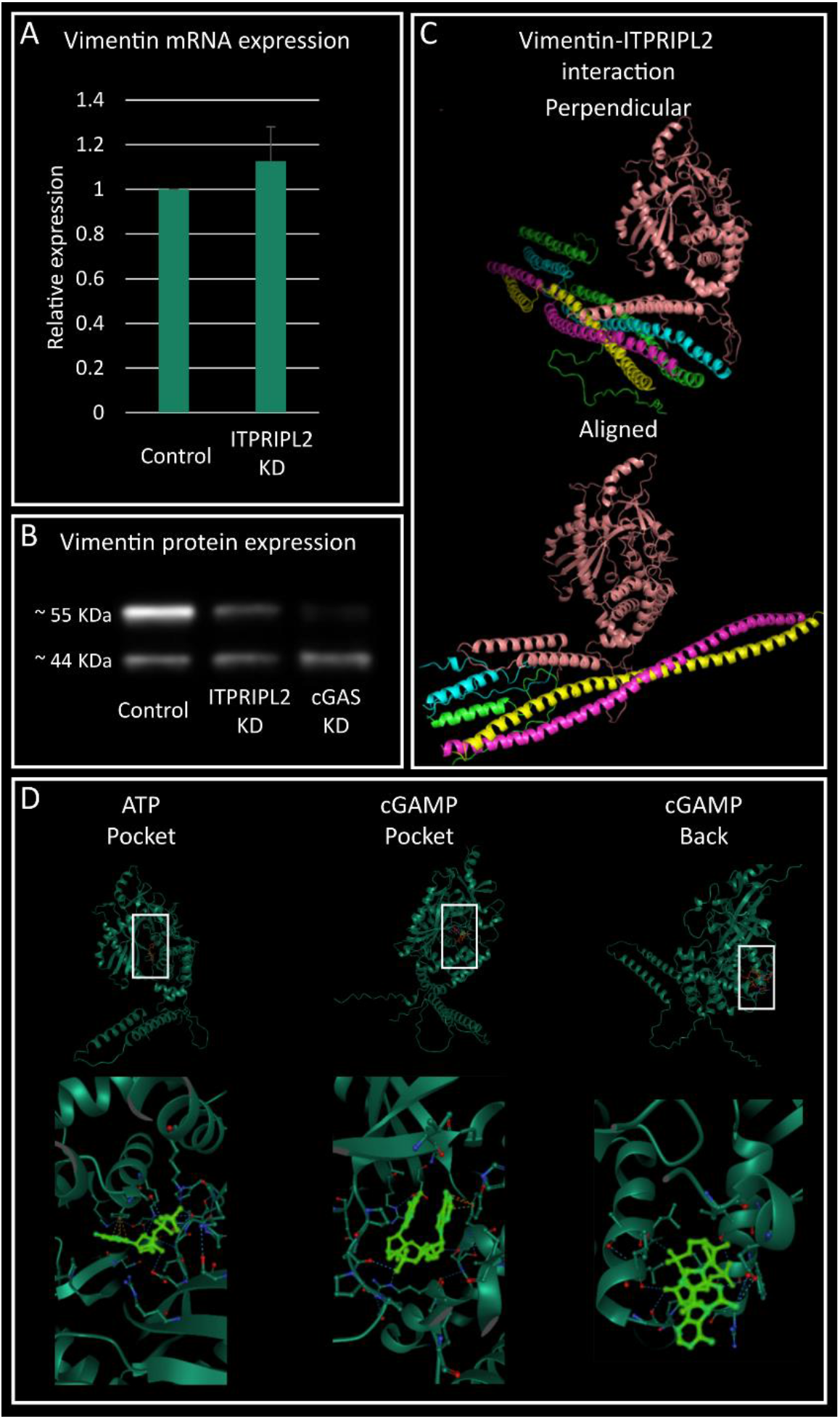
ITPRIPL2 is associated with vimentin. A) Vimentin mRNA levels were not altered upon ITPRIPL2 knockdown. B) Western blot analysis of vimentin in nHDF cells did not show a decrease in vimentin abundance upon ITPRIPL2 knockdown, however the ratio between the two variants shifted from 6:1 to 2:1, indicating an alteration of the vimentin protein in absence of ITPRIPL2. Knockdown of cGAS, the enzyme responsible for cGAMP production, resulted in an even more severe shift in vimentin form, to a 0.4:1 ratio. C) Protein docking simulation of vimentin and ITPRIPL2 with pyDockWEB identified two potential interaction configurations with favourable binding energy. For one possible configuration, the horseshoe domain of ITPRIPL2 binds perpendicular to the vimentin strand and for the other configuration ITPRIPL2 binds at the transition between vimentin monomers. D) Assessment of potentially missing ligands with AlphaFill identified two potentially relevant ligands, ATP and cGAMP. Both molecules could bind in the central ITPRIPL2 pocket of ITPRIPL2. Additionally, a secondary potential binding site for cGAMP was identified at the back of ITPRIPL2.

### 2.4 Protein interaction modelling of Vimentin and ITPRIPL2

To explore a potential interaction between vimentin and ITPRIPL2, a protein docking simulation was performed. For vimentin a recently published tetramer structure (elongated state) was used [36], as this state most likely reflects the conformation present in intermediary filaments. For ITPRIPL2 the AlphaFold2 [37] predicted structure obtained from AFDB was used [38, 39]. We opted for pyDockWEB to assess potential interaction between these proteins. PyDockWEB was ran with binding constraints for the residues in the horseshoe domain as these domains are known to be involved in protein-protein interactions [40]. The results of the docking simulation suggest two potential interaction configurations: one where the horseshoe domain of ITPRIPL2 binds perpendicular across the vimentin strands or an alternative configuration where it binds at the transition between different monomers that make up the vimentin tetramer (Figure 4C). The docking simulation cannot conclusively determine which is the more likely configuration but both have favourable (negative) binding energy (Table S2).

This model was further refined by assessing potentially missing ligands with AlphaFill [41]. Only low identity results (25%) were available for ITPRIPL2. Two potentially relevant ligands are ATP and cGAMP. Both molecules could bind in the central pocket of ITPRIPL2 (Figure 4E). Further, there is a secondary potential binding site for cGAMP at the back of ITPRIPL2 (opposite side of the horseshoe domain, Figure 4D). To determine whether cGAMP can serve as a co-factor for ITPRIPL2, we assessed the effect of cGAS (the enzyme that produces cGAMP) knockdown on vimentin. Western blot analysis of cGAS knockdown cells resulted in a similar shift in vimentin type as present in case of ITPRIPL2 knockdown (Figure 4B). This supports the modelling results and shows that cGAMP is required for vimentin processing, likely through binding to ITPRIPL2.

### 2.5 Transcriptomics analysis

To determine the effect of ITPRIPL2 knockdown at a process level, transcriptomics data was generated from control and ITPRIPL2 knockdown in both HeLa and nHDF cells. Differential gene expression analysis showed a contrasting difference in the effect of ITPRIPL2 knockdown between the two cell types. In fibroblasts, over 1300 genes were differentially expressed, with 1142 downregulated and 245 upregulated genes compared to the control, while fewer differentially expressed genes were present in HeLa cells, 308 of which 54 were downregulated and 250 were upregulated genes (Figure 5A). No distinct changes were observed in the expression of known fission or fusion regulators in either cell type, while ITPRIPL2 expression was significantly decreased (Figure 5B). This implies that the effect of ITPRIPL2 knockdown on mitochondrial dynamics is not through manipulation of expression levels of the established mitochondrial dynamics proteins.

**Figure 5:**
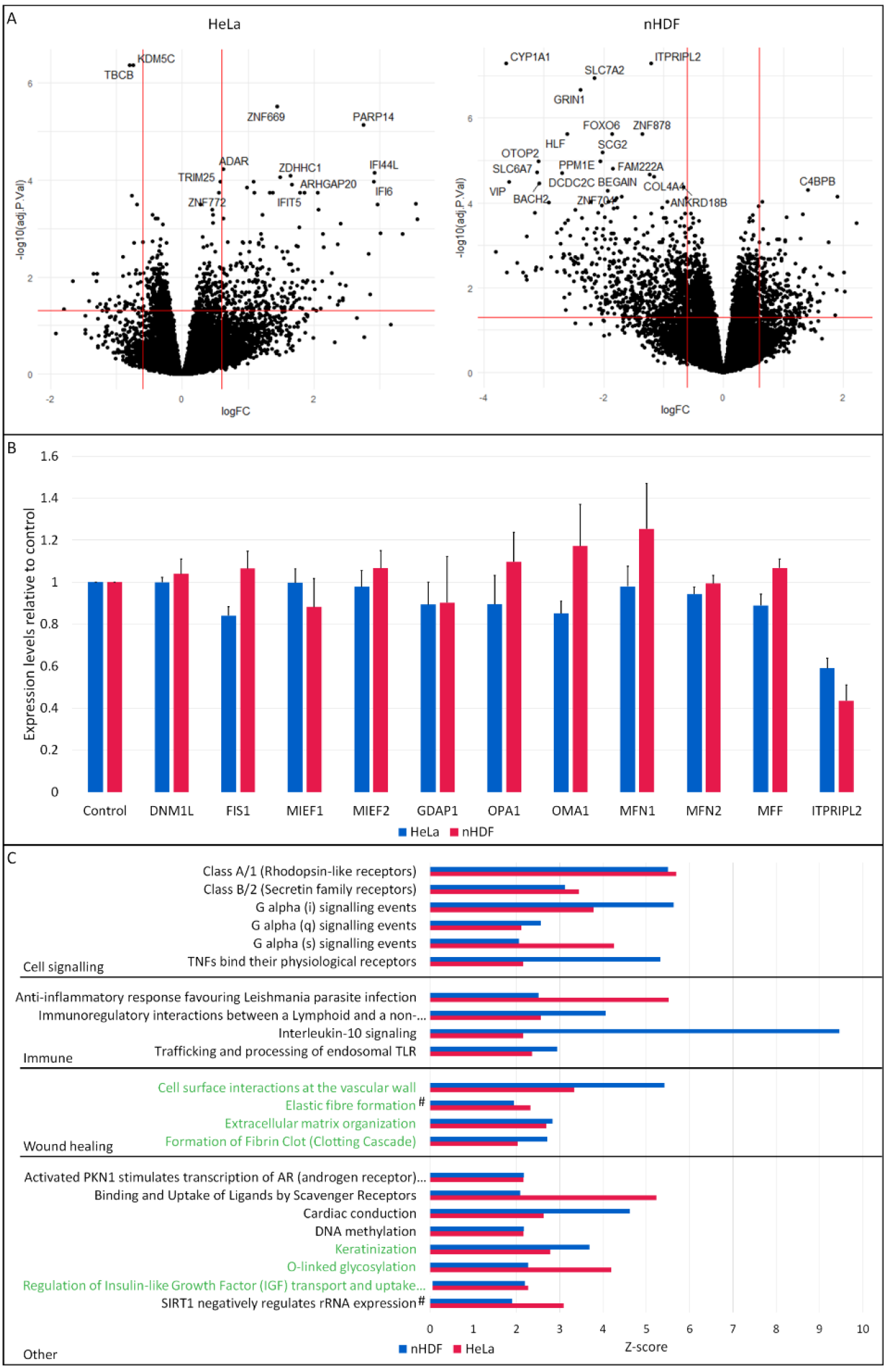
Transcriptomics analysis of ITPRIPL2 knockdown compared to non-targeting (GFP) knockdown in HeLa cells (N=3) and nHDF (N=4). A) Distribution of differentially expressed genes (DEGs) shows a different pattern in HeLa and nHDF cells, with 308 DEGs in HeLa, most of which are upregulated genes, whereas over 1300 DEGs were found for nHDF cells, with mostly downregulated genes. B) Relative expression of fission and fusion regulators upon ITPRIPL2 knockdown showed no distinct changes compared to control cells in either cell type, while the expression of ITPRIPL2 is 40-50% decreased. C) Pathways significantly enriched in both cell types (Z-score≥1.96). Enriched pathways can be categorised into cell signalling, immune related, wound healing and other pathways. Seven pathways with an established link to vimentin were identified, indicated in green. Pathways indicated with # were significant for nHDF cells and approaching significance (Z-score≥1.9) in HeLa cells.

Pathway enrichment analysis showed 51 significantly enriched pathways in nHDFs and 58 in HeLa cells. However, none of the enriched pathways were directly linked to mitochondrial dynamics. Since the observed effect on mitochondrial morphology was the same in both cell types, we focussed on processes that were changed in both nHDFs and HeLa cells. Twenty pathways were significantly enrichment in both cell types and two pathways which were enriched in one and approaching significance in the other cell type. Ten of these pathways are related to either cell signalling or immune related processes (Figure 5C). Four pathways, ‘formation of fibrin clot (clotting cascade)’, ‘extracellular matrix organization’, ‘elastic fibre formation’ and ‘cell surface interactions at the vascular wall’ have a connection with wound healing, a process in which vimentin is involved. Three additional enriched pathways can be regulated by vimentin or are essential for proper vimentin functioning (Figure 5C). This data further supports a role for ITPRIPL2 in vimentin driven intermediate filament formation.

### 2.6 Vimentin knockdown results in mitochondrial elongation

Since ITPRIPL2 is an intermediate filament associated protein that influences mitochondrial dynamics, the effect of intermediate filament knockdown on mitochondrial morphology was assessed. Vimentin knockdown resulted in mitochondrial elongation (Figure 6A/B), similarly to ITPRIPL2, indicating that intermediate filaments are important for mitochondrial morphology.

**Figure 6:**
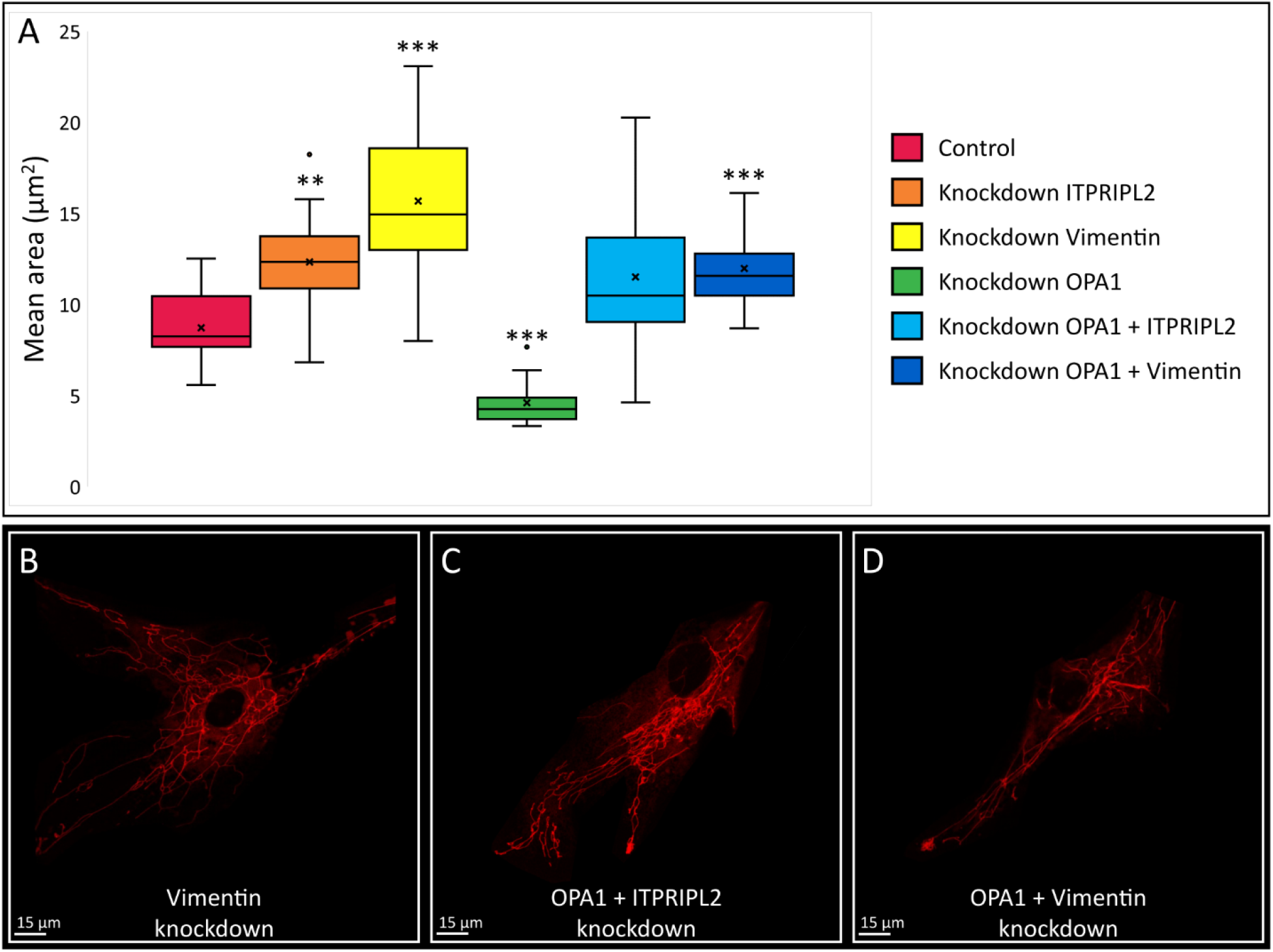
Assessment of mitochondrial morphology upon intermediate filament knockdown. A) Distribution of the mean mitochondrial area for control and knockdown conditions in human fibroblasts. A significant increase in mean area was observed for knockdown of ITPRIPL2, vimentin and OPA1+vimentin. For OPA1 knockdown a significant decrease in mean area was observed, while mitochondrial morphology was unchanged for the OPA1+ITPRIPL2 knockdown. B-D) Examples of mitochondrial network morphology for vimentin, OPA1+ITPRIPL2 and OPA1+vimentin knockdown conditions. Data information: Scale bar indicates 15 µm. Knockdown ITPRIPL2 N=11, Knockdown OPA1+ITPRIPL2 N=12, Control and Knockdown OPA1+vimentin N=13, Knockdown OPA1 N=14, Knockdown vimentin N=15. **p < 0.01, ***p < 0.001 (Mann Whitney U Test, significance level adjusted for multiple testing).

To assess whether the observed change in morphology was the result of a decrease in fission or an increase of fusion, double knockdowns were induced of OPA1 and ITPRIPL2 or vimentin. While OPA1 knockdown resulted in extensive mitochondrial fragmentation, the combination of OPA1 with ITPRIPL2 or vimentin knockdown revealed a normal to elongated mitochondrial network (Figure 6A/C/D). These results indicate that disruption of the intermediate filaments results in decreased levels fission instead of increased fusion.

## 3 Discussion

Previous studies have often identified proteins related to mitochondrial dynamics through diseases or protein homology [23, 30, 42]. However, these are time consuming efforts that result in the identification of one or two proteins at a time. To create a better understanding of the mitochondrial proteome, integrative large scale computational or experimental strategies are required [43, 44]. In this study, we employed a computational protein-protein interaction network approach to identify novel proteins or processes related to mitochondrial dynamics, based on the principle that proteins with a similar function or a function in the same process are more likely to interact with each other [45]. This computational approach to identify candidate proteins uses already available knowledge without the need for large-scale and costly experiments. Protein-protein interactions have been successfully applied to identify new functions for proteins and with the increase in available knowledge these predictions are becoming more accurate [43]. Additionally, the integration of multiple evidence types studying these interactions enables the identification of orphan proteins in relation to a process of interest.

### 3.1 Identification of candidates

The protein-protein interaction network was constructed based on established mitochondrial dynamics proteins and expanded with proteins based on their interaction with these established proteins. The number of expanded proteins and the confidence cut-off were selected carefully. Optimal values depend on the selected query proteins as well as currently available knowledge of the process of interest and should be optimized for every study individually. The confidence score determines the amount and type of evidence required for an interaction to be added to the network. A too high score can restrict the discovery of new information, while a too low confidence cut-off increases the risk of false positives. In exploratory research especially the confidence score cannot be too strict as there is limited information available for novel findings. Here, a medium confidence score of 0.5 was chosen, as it resulted in interactions based on a combination of text-mining, experimental, co-expression, and curated knowledge evidence types, with a good connectivity of the query proteins with the expanded proteins in the final network. Importantly, the network also contained multiple proteins without known function, providing interesting insight into their potential function.

Leave-one-out validation revealed that 72% the query proteins were present in the network upon expansion, as would be expected based on the principle underlying the network approach highlighting a robust final network. However, a few query proteins did not re-occur in the network, including PLD6, STOML2, MTFR1, MSTO1 and SLC25A46. More research into these proteins and their involvement in biological processes could result in sufficient evidence to connect them to the others. Furthermore, the list of candidates contained a significant overrepresentation of mitochondrial proteins and multiple proteins previously associated with mitochondrial dynamics, with nine proteins amongst the twenty best connected proteins, including INF2 and OMA1 [3, 46]. Their presence in the network combined with the overrepresentation of mitochondrial proteins supports the validity of this analysis for identifying novel mitochondrial dynamics proteins.

### 3.2 Functional validation of candidate genes

Four candidates without a known function were selected based on their heat propagation ranking. To determine whether these candidates influenced mitochondrial dynamics, gene knockdowns were created and changes in mitochondrial morphology were assessed. Knockdown of all four selected candidates showed an increased mean mitochondrial area and network complexity (Figure 2), which implies a misbalance between fission and fusion. Two of these candidates, WBP1L and MB21D2, have interactions with the known Drp1 adaptors, MIEF1 and MIEF2, while WBP1L also interacts with MFF. While it is currently uncertain how these proteins affect mitochondrial dynamics precisely, the interactions with known Drp1 adapters suggest that these proteins could influence the recruitment of Drp1. For TMEM126A, a function in complex I assembly was established after candidate selection [47, 48]. Although knockdown of TMEM126A resulted in an elongated mitochondrial network, there is no clear connection between complex I and mitochondrial dynamics. However, a shift in the balance towards mitochondrial fusion can occur upon stress to increase energy production [9]. Impaired complex I assembly and functioning is linked to oxidative stress [49], implying that the observed effect on mitochondrial dynamics is indirect.

The role of ITPRIPL2 in mitochondrial dynamics was investigated in more detail, since knockdown of this protein resulted in the most severe mitochondrial morphology change. It was initially expected that ITPRIPL2 would localize to the mitochondria, or an organelle which is linked to mitochondrial dynamics, like the ER. Interestingly, our data shows that ITPRIPL2 co-localises with a type III intermediate filament protein, vimentin (Figure 3). This association was confirmed by changes in the shape and organisation of vimentin positive intermediate filaments upon ITPRIPL2 knockdown and overexpression (Figure 4). The decrease in vimentin staining upon ITPRIPL2 knockdown is not the result of an absolute decrease in presence of vimentin, as was shown by western blot analysis, but rather a shift between the two observed vimentin forms (Figure 4B). Although vimentin has one isoform of around 55-57 kDa in size according to the RefSeq database [50], we observed a second form of approximately 44 kDa in size. Multiple protein forms of vimentin have been observed on western blot with different antibodies (Santa Cruz Biotechnology, sc-66002; Progen, 64001; Proteintech, 10366-1-AP; Bioss, bs-0756R; Abcam, ab45939; [51, 52]), including the 44 kDa form. Moreover, cleavage of vimentin has been reported upon induction of apoptosis, resulting in 44, 36, 25 and 15 kDa peptides [53]. It is however unlikely that the observed form of 44 kDa is the result of apoptosis induced cleavage as the other peptides were not observed in our experiments. The abundant presence of the smaller vimentin form in knockdown cells suggests a processing step, for instance the addition of post-translational modifications, which is inhibited by ITPRIPL2 knockdown. The occurrence of a processing step before or during assembly of vimentin for which ITPRIPL2 is essential, would be in line with the absence of assembled vimentin structures in the immunostaining upon ITPRIPL2 knockdown.

Docking simulations of ITPRIPL2 and vimentin were done to assess the potential interaction between ITPRIPL2 and vimentin, which suggested two potential interaction configurations. While the more likely configuration could not be determined, both have a favorable binding energy, suggesting that docking between ITPRIPL2 and vimentin is likely. Alphafill predictions indicated possible missing ligands and suggested ATP and cGAMP as potentially relevant ligands. As vimentin levels have previously been linked to cGAMP treatment [54], it was further validated as a potential ITPRIPL2 ligand. Knockdown of cGAS, the enzyme that produces cGAMP, resulted in an even more extreme shift between vimentin forms (Figure 4B). Additionally, knockdown of cGAS results in mitochondrial elongation, similar to ITPRIPL2 knockdown (Data not shown). This shows that cGAMP is a likely ligand necessary for proper ITPRIPL2 functioning.

### 3.3 Transcriptomics analysis

To gain further insight into the cellular function of ITPRIPL2, transcriptomics data was generated for ITPRIPL2 knockdown in both HeLa and nHDF cells. These two cell types display a similar effect on mitochondrial morphology, indicating the same underlying mechanism leading to the change in mitochondrial morphology. The distribution of differentially expressed genes was distinctly different between the two cell types, emphasizing the importance of using multiple cell types to exclude cell type specific changes. The transcriptomic data did not reveal any changes in expression of known fission or fusion genes in either cell type (Figure 5B) indicating that the observed effect on mitochondrial dynamics is not mediated through modulation of established mitochondrial dynamics proteins at the transcriptomics level but via a different mechanism.

Pathway enrichment analysis showed that half of the enriched pathways were linked to either immune response or the cell signalling, which is likely an effect of the change in mitochondrial morphology.

Imbalanced fission and fusion leads to mitochondrial dysfunction, which can activate early-phase inflammatory mediators and lead to the production of pro-inflammatory cytokines [55]. Furthermore, mitochondria are known to be involved in retrograde signalling, for instance mitochondrial calcium interacts with intracellular signalling pathways and mitochondrial ROS can block cell cycle progression [56, 57]. While the exact role of mitochondrial morphology remains unknown, transitions in morphology and positioning likely plays an important role in these signalling events [56, 57].

Interestingly, amongst the other pathways, several links to vimentin were observed (Figure 5C), including enrichment of four pathways related to wound-healing. Vimentin translocates towards the extracellular matrix upon injury and coordinates multiple actions required for proper wound healing, like fibroblast proliferation and collagen accumulation [58, 59]. In various studies vimentin knockout mice were shown to have impaired wound healing [58, 60] and cells with a vimentin knockout inhibit fibroblast growth, which decreases the collagen production necessary for wound healing [58]. This fibroblast inhibition and decreased collagen production eventually results in decreased keratinocyte mobility and a loss of keratinisation [58]. Enrichment of the keratinisation pathway, with an overall downregulation following ITPRIPL2 knockdown, suggests that similar processes are affected upon both vimentin and ITPRIPL2 depletion.

Next to wound healing and keratinisation, a pathway on regulation of insulin-like growth factor 1 was significantly enriched upon ITPRIPL2 knockdown and showed an overall upregulation. Extracellularly translocated vimentin has been linked to the activation of insulin-like growth factor 1, again linking ITPRIPL2 and vimentin [61]. In addition, the O-linked glycosylation pathway was overrepresented amongst the differentially expressed genes. This post-translational modification is important for the interaction between vimentin molecules, which in turn is crucial for the proper assembly of vimentin filaments [62]. The upregulation of this pathway potentially indicates that changes in the post-translational modifications of vimentin occur upon ITPRIPL2 knockdown.

### 3.4 Intermediate filaments are important for mitochondrial fission

While mitochondria and intermediate filaments are known to interact with each other [63], the link between mitochondrial fission and fusion and intermediate filaments is not well established. Intermediate filaments are linked to mitochondrial motility [63, 64], although evidence on changes in fission and fusion balance upon knockdown of intermediate filaments is limited [65]. Where a previous study reported mitochondrial fragmentation in case of vimentin knockdown, our data shows extensive mitochondrial elongation upon knockdown of vimentin and a vimentin associated protein (Figure 6), pointing towards a different mechanism. Moreover, double knockdowns of vimentin and OPA1 showed that elongation is present, even with significantly decreased levels of fusion (Figure 6), indicating that elongation is the result of a decrease in fission, rather than an increase in fusion.

Overall, our data reveals that ITPRIPL2, a protein with a previously unknown function, is associated with vimentin positive intermediate filaments. Moreover, we show that maintenance of intermediate filament structure is essential for mitochondrial fission. The data presented here provide a novel link between intermediate filaments and mitochondrial fission expanding our understanding of mitochondrial dynamics and involved regulators. Additionally, this study demonstrates that a computational protein-protein interaction network approach can successfully be used to identify novel proteins in mitochondrial dynamics, with a potential use for other biological processes.

## 4 Materials and Methods

### 4.1 Computational candidate selection

A query list of known proteins involved in mitochondrial dynamics was constructed based on literature and used to create a protein-protein interaction network using the stringApp (version 1.5.0 [66, 67]) in Cytoscape (version 3.7.1) [68]. Subsequently, the set of query genes was expanded with a set number of proteins, based on the total connectivity to the query set, compared to their overall connectivity in the database [67]. Confidence scores were tested between medium (0.4) and high (0.7) to determine optimal settings for the final network. The network was validated by excluding one protein from the query list and determining how often the removed proteins were present in the expanded network. Moreover, to confirm whether an overrepresentation of mitochondrial proteins was present in the expanded network, a hypergeometric test was performed using the dhyper function in R (version 4.1.3). For this, the number of mitochondrial proteins was determined based on MitoCarta3.0. To determine the most promising candidates, a heat propagation analysis was performed using the Diffusion app (version 1.5.4) in Cytoscape, as an indication of ‘network closeness’ for each protein to the initial query set.

### 4.2 Transient transfections

An esiRNA-mediated knockdown was created for OPA1, DNM1L, WBP1L, TMEM126A, MB21D2, ITPRIPL2, VIM, cGAS and a non-targeting control (GFP). T7 promotor sequence containing primers were used to amplify around 600bp of the respective transcripts. dsRNA was synthesized using the MEGAscript Kit (Ambion, AM1334) according to manufacturer protocol. For esiRNA production, dsRNA was incubated with Shortcut RNase III (NEB, M0245S) according to manufacturer instructions, with 0.75 units of enzyme per 1 µg dsRNA. To induce knockdown cells were transfected with esiRNAs for 48 to 72 hours. In short, esiRNAs were combined with RNAiMAX (Invitrogen, 13778075) in OptiMEM medium (Gibco, 11058021) with a concentration of 50 ng esiRNA/ml culture medium. Further experiments were performed 48 or 72 hours after transfection.

For overexpression, 1 µg plasmid was combined with 3 µl FuGENE (Promega, E2311) in OptiMEM medium for each ml of culture medium. Following a 30 minute incubation at room temperature, cells were transfected with the plasmid/FuGENE mixture and incubated for 24 to 48 hours before followed up experiments were performed.

### 4.3 Knockdown efficiency

Normal Human Dermal Fibroblasts (nHDF) seeded in a 6-well plate at a density of 10,000 cells per well respectively were transfected with esiRNA. RNA was isolated 72h post esiRNA transfection with the RNeasy Mini Kit (Qiagen, 74104) and the RNAse-Free DNase Set (Qiagen, 79254) according to manufacturer instructions. cDNA was synthesized with the qScript cDNA Supermix kit (Quantabio, 95048- 500), using 400 ng RNA as input. qPCR was performed using 1x SensiMixTM SYBR® (Bioline Meridian, QT615-05 2x stock) and a final primer concentration of 0.5 μM, on the LightCycler® 480 II (Roche, 0501524300). Knockdown efficiency was determined by calculating the ΔΔCt between knockdown and control samples, using *TBP* for normalization. Significance of the relative differences was determined using a Mann Whitney U Test (R version 4.1.3 and RStudio build 461).

### 4.4 Mitochondrial morphology

nHDF cells were seeded onto 8 well µ-Slides (Ibidi, 80826), with a density of 1000 cells. Upon 48 and 72h of knockdown mitochondria were stained with MitoTracker Red FM (300nM, ThermoFisher Scientific, M22425). MitoTracker containing medium was replaced with imaging medium of DMEM without phenol red (Gibco, 21063029). Randomly selected cells were imaged at room temperature using a laser scanning confocal microscope (Leica DMI4000B microscope with a Leica TCS SPE confocal system), with a 63x oil objective (numerical aperture 1.3) and the Leica LAS-AF acquisition software. Deconvolution was applied to all images using the Iterative Deconvolve 3D plugin, without a wiener filter gamma and with a low pass filters of 1, a maximum number of iterations of 5 and a termination of iterations at 0.010 [69]. Mitochondrial morphology was assessed using the Mitochondria Analyzer plugin for FIJI/ImageJ [70]. Thresholding settings were adjusted, with enhance local contrast turned off, a block size of 10 and a C-value of 2, as previously assessed for nHDFs, before analysis with default settings [71]. Per condition, 11- 18 nHDF cells were analyzed and statistical significance was determined using Mann Whitney U Tests, adjusted for multiple testing using Bonferroni correction (R version 4.1.3 and RStudio build 461). All quantifications were verified in multiple independent experiments.

### 4.5 Immunostaining

nHDF or HeLa cells were seeded onto chamber slides (ThermoFisher, 177429) with a density of 10.000 or 25.000 cells respectively. Upon knockdown or overexpression, slides were fixed in 3.7% formaldehyde, permeabilized in 0.1% PBS-Tween-20 and blocked with 3% BSA. Primary antibodies against vimentin (Proteintech, 10366-1-AP 1:100), tom20 (BD biosciences, 612278 1:100) and FLAG (Sigma, A9594 1:500) were diluted in 3% BSA and incubated for 1 hour at room temperature, followed by a 1 hour incubation with 1:1000 diluted secondary antibodies (Invitrogen A11034 and A21424). Slides were airdried, mounted with ProLong Gold Antifade reagent (Invitrogen, P36941/P10144) and imaged at room temperature using a laser scanning confocal microscope (Leica DMI4000B microscope with a Leica TCS SPE confocal system), with a 63x oil objective (numerical aperture 1.3) and the Leica LAS-AF acquisition software.

### 4.6 Western blot

Whole cell protein was extracted from nHDF cells with lysis buffer containing 50mM Tris, 150 mM NaCl, 1 mM EDTA, 0.01% Triton X-100 and 1x protease inhibitor. Per sample, 20 µg protein was separated with SDS-page (Bio-Rad, 4561084DC) and blotted on a PVDF membrane (Bio-Rad, 1704156) using a Trans-Blot Turbo Transfer System (Bio-Rad, 1704150). Membranes were blocked with 5% non-fat dry milk and incubated overnight at 4 °C with primary antibodies against vimentin (1:5000), followed by a 1 hour incubation at room temperature with secondary antibodies (Santa Cruz, sc-2004 1:5000). Chemiluminescence (Santa Cruz, sc-2048) was used for visualisation of proteins on a ChemiDoc XRS imaging system (Bio-Rad).

### 4.7 Protein docking simulations

Vimentin tetramer structures were obtained from the authors of Vermeire et al 2023 [36]. The predicted structure of ITPRIPL2 was obtained from AFDB (UniProt identifier Q3MIP1). Protein-protein interactions were assessed using pyDockWEB (pyDock3 version 3.5.1) [72]. Due to limitations in the configuration of pyDockWEB, input molecules are size limited (∼160 Å diameter). We therefore used PyMOL to cut the vimentin tetramer into 4 roughly equal sized, slightly overlapping chunks along the length axis of the tetramer. The four sections correspond to: a 150 Å diameter sphere from the most distal residues on each end of the tetramer (residue 410 on chain A and residue 400 on chain D for chunks 1 and 4 respectively) and 2 spherical volumes with a radius of 75 Å, the first around residue 225 on chain A (chunk 2) and the second around residue 139 on chain A (chunk 3). The ITPRIPL2 structure was aligned to each chunk with the outward (“down”) facing residues of the horseshoe domain selected as constraints (see Table S3 for selected residues). Supplementary Table S2 shows the best configurations achieved for each chunk. The configurations where both constrained and unconstrained energies were highest ranked, were considered the most likely docking positions.

We used the AlphaFill public server (v2.1.0) to determine potential ligands. The best candidate positions for ATP and cGMAP were optimised with the built-in call to Yasara to minimise the transplant crash score. Supplementary table S4 shows the initial scores and optimised scores for selected positions.

### 4.8 Transcriptomics analysis

Knockdown of ITPRIPL2 and GFP were induced in nHDF (N=4 for each condition) and HeLa cells (N=4 for each condition) and RNA was isolated with the RNeasy Mini kit (Qiagen, 74104) including DNAse treatment, following manufacturers instructions. RNA was enriched for mRNA using Nex Poly(A) Beads 2.0 (Perkin Elmer, NOVA-512992). Library preparation was performed with the Rapid Directional RNA-Seq Kit 2.0 (Perkin Elmer, NOVA-5198-02) with NEXTFLEX Unique Dual Index Barcodes. Sequencing data was generated with a NovaSeq 6000 (Illumina) on a single NovaSeq 200 cycle S1 flowcell.

Reads were processed and mapped to the HG38 reference genome using RNA STAR on the Galaxy platform [73, 74]. Aligned reads were converted to transcript counts on the Galaxy platform. In our desire to focus our analyses on protein-coding genes, genes with less than 10 in more than 70% of samples were not considered for downstream analysis. Counts were CPM normalized and log2 transformed using the limma package in R (R version 4.1.3 and RStudio build 461)[75]. Batch effects were first assessed using PCA analysis. For the HeLa samples, one sample was stored several months longer than the other three before sequencing. PCA analysis showed a clear separation between this sample and the other replicates and was hence not considered for further analysis. This resulted in N=3 for both ITPRIPL2 and GFP knockdown HeLa samples. Batch effects for the remaining samples were removed by applying the removeBatchEffect function in limma [76].

Differentially expressed genes (DEGs) were identified between ITPRIPL2 and non-targeted GFP knockdown data using the limma package in R, by creating a full design matrix with knockdown status and cell type as co-variates and calculating the desired contrast. Genes were considered to be differentially expressed with an absolute log2 fold change of 0.58 (equal to a fold change of 1.5) and an uncorrected p-value ≤ 0.05. Using the uncorrected p-value is warranted by the focus on pathway enrichment analysis, which mitigates part of the multiple testing problem [77]. Pathway analysis was performed in PathVisio [78], with the REACTOME pathways hs_derby_ensemble_103.bridge as a gene database. Pathways were considered enriched when having a Z-score ≥1.96 and containing at least 3 DEGs.

### 4.9 Data availability

Scripts used for the creation and analysis of the networks can be found on GitHub at https://github.com/IreneHemel/MitochondrialDynamics.

The RNA sequencing data described in this manuscript have been deposited to the GEO database (https://www.ncbi.nlm.nih.gov/geo/) and assigned the identifier GSE232284.

## Supplemental material

Document S1. Figures S1–S2 and Table S1-S4.

## Author contributions

**Irene Hemel:** Investigation, Software, Validation, Formal analysis, Data Curation, Visualization, Writing - Original Draft, Writing - Review & Editing. **Carlijn Steen:** Investigation, Software, Validation, Formal analysis. **Simon Denil:** Formal analysis, Writing – Review & Editing. **Gökhan Ertaylan:** Writing - Review & Editing, Supervision. **Martina Kutmon:** Methodology, Writing – Review & Editing. **Michiel Adriaens:** Methodology, Software, Writing – Review & Editing. **Mike Gerards:** Conceptualization, Resources, Writing - Review & Editing, Supervision.

