## Supplemental information for "The guardians of mitochondrial dynamics: a novel role for intermediate filament proteins"

### Supplement

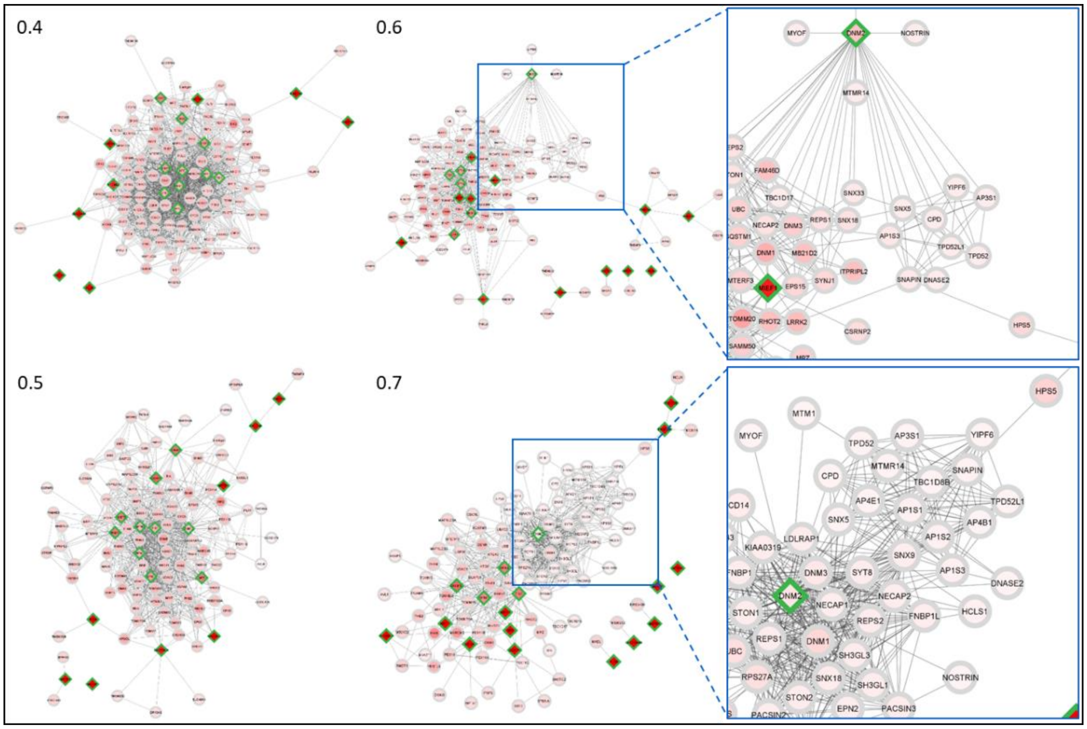


*Figure S1: Comparison of protein-protein interaction networks created with different cut-offs. Networks with 100 expanded proteins and differing confidence cut-off scores. Confidence scores above 0.5 result in the formation of subclusters within the network, with proteins mostly interacting with each other, instead of with the query set, while a low score results in the inclusion of proteins and connections for which little evidence is present.*

*Table S1: List of candidates with a previous association with mitochondrial dynamics.*

| Gene | Rank | Function in dynamics | Reference |
|---|---|---|---|
| **INF2** | 1 | Actin polymerization at mitochondrial-ER contact sites, resulting in mitochondrial pre-constriction. | [1, 2] |
| **YME1L1** | 3 | Proteolytic cleavage of OPA1. Knockdown leads to elongation of mitochondria. | [3, 4] |
| **OMA1** | 4 | Proteolytic cleavage of OPA1. | [5] |
| **MARCH5** | 7 | Knockdown can result in mitochondrial elongation or fragmentation. Possible ubiquitination of FIS1, DNM1L and MFN1. The RING-domain is important for proper functioning. | [6-9] |
| **PHB2** | 9 | Regulates activity of SLP2 (*STOML2)* and the ratio of long-OPA1 to short-OPA1. | [10] |
| **MUL1** | 10 | Degradation of Mfn1/2. | [11] |
| **PARL** | 11 | Cleaves PGAM5 and PINK1 and expression of cleaved PARL induces fragmentation. | [12, 13] |
| **AFG3L2** | 13 | Regulation of OMA1 mediated cleavage of OPA1. | [14] |
| **PINK1** | 20 | Mitophagy, activation of Drp1 and promotes degradation of Mfn1/2. | [11, 15-17] |
| **IMMT** | 24 | Knockdown results in decreased fusion and fission activity trough decreased levels of OPA1, Mfn1/2, Drp1, Mid49 and MFF. | [18] |
| **OPA3** | 27 | Regulation of fission, likely independent of Drp1 and FIS1. | [19] |
| **PRKN** | 30 | Mitophagy and degradation of Mfn1/2. | [11, 15, 17] |
| **SPG7** | 42 | Mutations leading to increased expression result in fragmented mitochondria. | [20] |
| **PGAM5** | 43 | Mitophagy and activation/recruitment of Drp1. | [21, 22] |
| **SH3GLB1** | 47 | Knockdown results in abnormally interconnected mitochondria dependent on the presence of Drp1. | [23] |

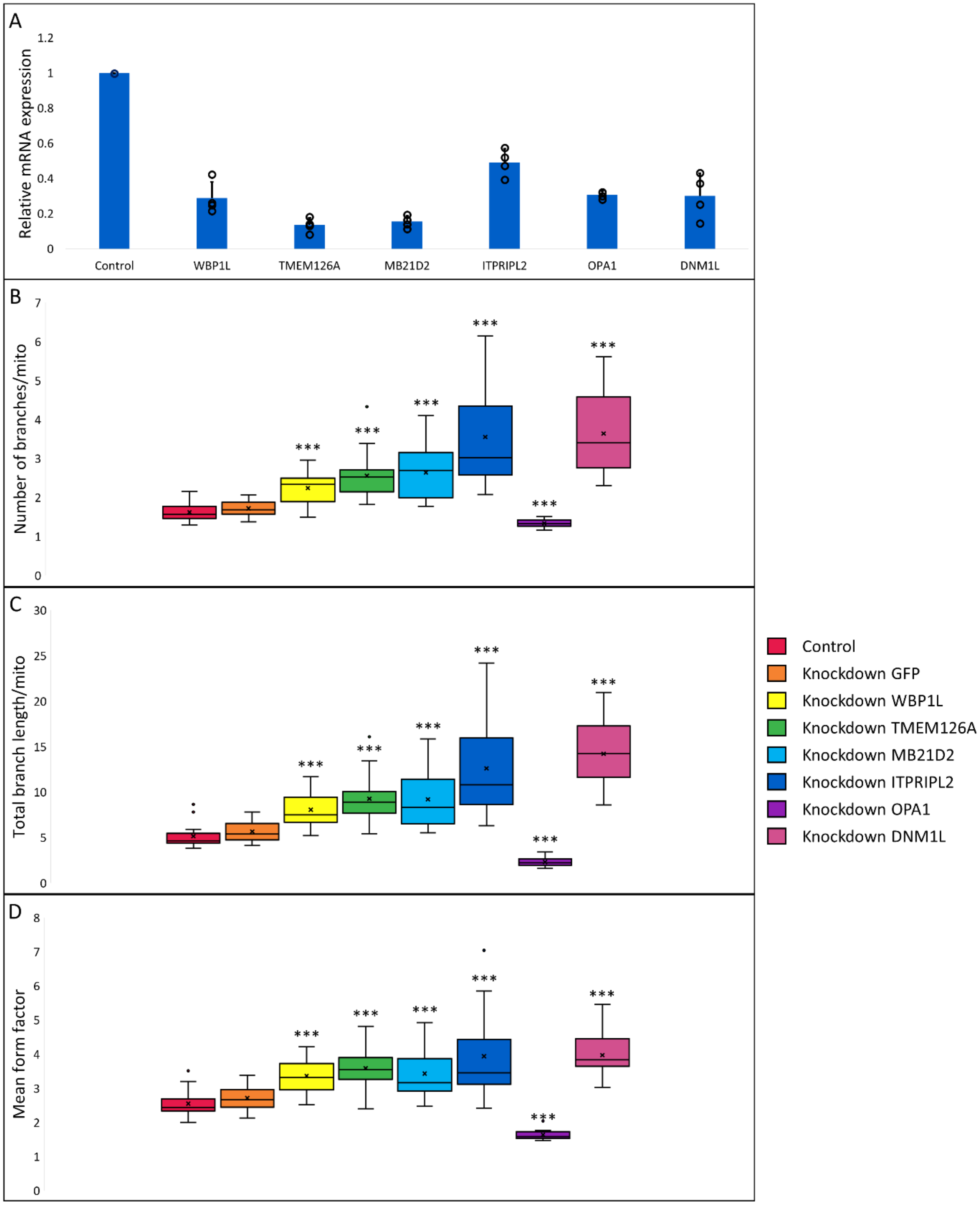


*Figure S2: Assessment of knockdown efficiency and mitochondrial morphology.*

*A) All knockdown conditions show at least a 50 percent decrease in mRNA expression in normal human dermal fibroblasts (nHDF). Error bars indicate s.d., N=4 for all conditions.*

*B-D) Distribution of parameters indicative of network complexity provided by the Mitochondrial Analyzer for control and candidate knockdown conditions in normal human dermal fibroblasts.*

*B) The number of branches per mitochondrion was significantly increased for all candidate knockdowns, indicating an increase in mitochondrial network complexity. For OPA1 and DNM1L a respective decrease and increase in the number of branches was found, while no changes were present upon GFP knockdown.*

*C) A significant increase in the total branch length per mitochondrion was observed in case of WBP1L, TMEM126A, MB21D2 and ITPRIPL2 knockdown. For positive controls OPA1 and DNM1L a significant decrease and increase in total branch length was found, respectively, while branch length was unchanged for negative control GFP*
*D) The mean form factor, a parameter indicative of mitochondrial shape was significantly increased for all candidate knockdowns, indicating an increase in mitochondrial network complexity. For OPA1 and DNM1L a respective decrease and increase in the mean form factor was found, while no changes were present upon GFP knockdown.*
*Data information: Control N=18, Knockdown GFP, MB21D2 N=16, Knockdown WBP1L, TMEM126A, ITPRIPL2, OPA1 N=15, Knockdown DNM1L N=13. ***p < 0.001 (Mann Whitney U Test, significance level adjusted for multiple testing). Experiment was repeated 4 times.*

*Table S2: Docking simulation configurations for each vimentin chunk, for both constrained (EneRST) and unconstrained (Ene) conditions. Selected possible configurations are highlighted.*

| | EneRST | | | | | | Ene | | | | | |
|---|---|---|---|---|---|---|---|---|---|---|---|---|
| | **Conf** | **Ele** | **Desolv** | **VDW** | **relRST** | **Total** | **Conf** | **Ele** | **Desolv** | **VDW** | **Total** | **RANK** |
| ***Chunk 1*** | 380 | -22.3 | -31.2 | 85.5 | 93.8 | -138.7 | 3675 | -47.8 | -17.8 | 26.2 | -62.9 | 1 |
| | 1394 | -22.2 | -27.8 | 51.7 | 93.8 | -138.6 | 7221 | -50.5 | -14.5 | 30.1 | -62.0 | 2 |
| | 6573 | -42.6 | -20.8 | 33.3 | 75.0 | -135.1 | 6573 | -42.6 | -20.8 | 33.3 | -60.1 | 3 |
| | 3433 | -40.0 | -16.2 | 30.8 | 81.3 | -134.4 | 3491 | -59.2 | -4.8 | 40.9 | -59.9 | 4 |
| | 3194 | -47.8 | -15.4 | 45.1 | 75.0 | -133.7 | 3194 | -47.8 | -15.4 | 45.1 | -58.7 | 5 |
| | 2148 | -28.2 | -34.6 | 101.1 | 75.0 | -127.7 | 9498 | -43.0 | -27.9 | 122.6 | -58.6 | 6 |
| | 9498 | -43.0 | -27.9 | 122.6 | 68.8 | -127.4 | 7380 | -40.6 | -18.9 | 12.8 | -58.2 | 7 |
| | 95 | -28.3 | -21.1 | 104.8 | 87.5 | -126.4 | 5390 | -46.1 | -16.8 | 48.2 | -58.1 | 8 |
| | 2915 | -48.5 | -13.3 | 43.0 | 68.8 | -126.2 | 2915 | -48.5 | -13.3 | 43.0 | -57.5 | 9 |
| | 818 | -33.0 | -11.8 | 81.2 | 87.5 | -124.2 | 7908 | -47.7 | -16.8 | 78.2 | -56.6 | 10 |
| | 5785 | -36.1 | -12.8 | 45.2 | 75.0 | -119.4 | 9439 | -54.4 | -7.7 | 74.5 | -54.6 | 11 |
| | 7221 | -50.5 | -14.5 | 30.1 | 56.3 | -118.2 | 8419 | -41.8 | -13.2 | 6.4 | -54.4 | 12 |
| | 7248 | -34.1 | -12.6 | 36.1 | 75.0 | -118.2 | 3744 | -41.1 | -19.5 | 63.9 | -54.2 | 13 |
| ***Chunk 2*** | 5180 | -51.4 | -14.7 | 52.4 | 87.5 | -148.3 | 6557 | -54.3 | -25.5 | 32.5 | -76.5 | 1 |
| | 5424 | -55.6 | -29.2 | 126.5 | 75.0 | -147.1 | **2904** | **-52.9** | **-26.8** | **42.4** | **-75.5** | **2** |
| | **2904** | **-52.9** | **-26.8** | **42.4** | **68.8** | **-144.2** | 5424 | -55.6 | -29.2 | 126.5 | -72.1 | 3 |
| | 5496 | -51.9 | -14.9 | 105.1 | 87.5 | -143.8 | 6989 | -47.4 | -25.7 | 31.2 | -69.9 | 4 |
| | 6525 | -61.2 | -5.1 | 3.7 | 75.0 | -140.9 | 7997 | -49.6 | -27.5 | 90.9 | -68.0 | 5 |
| | 5495 | -37.1 | -34.3 | 62.2 | 75.0 | -140.1 | 9569 | -48.9 | -20.0 | 21.9 | -66.6 | 6 |
| | 4879 | -55.6 | -14.3 | 51.7 | 75.0 | -139.7 | 6525 | -61.2 | -5.1 | 3.7 | -65.9 | 7 |
| | 6557 | -54.3 | -25.5 | 32.5 | 62.5 | -139.0 | 2965 | -52.4 | -21.1 | 80.6 | -65.5 | 8 |
| | 3101 | -55.7 | -6.8 | 60.7 | 81.3 | -137.7 | 5898 | -44.1 | -23.9 | 25.2 | -65.5 | 9 |
| | 8185 | -34.4 | -26.4 | 61.5 | 81.3 | -135.9 | 5495 | -37.1 | -34.3 | 62.2 | -65.1 | 10 |
| | 8682 | -31.2 | -24.2 | 21.3 | 81.3 | -134.6 | 2102 | -68.3 | 4.0 | -6.6 | -65.0 | 11 |
| | 1237 | -42.7 | -25.7 | 66.6 | 68.8 | -130.5 | 5024 | -68.1 | 4.9 | -15.5 | -64.8 | 12 |
| | 2341 | -37.8 | -27.9 | 45.8 | 68.8 | -129.8 | 4879 | -55.6 | -14.3 | 51.7 | -64.7 | 13 |
| ***Chunk 3*** | 5015 | -40.2 | -21.2 | 58.8 | 50.0 | -105.5 | 80 | -66.6 | 7.6 | -32.7 | -62.3 | 1 |
| | 5494 | -27.9 | -20.8 | 66.8 | 51.9 | -94.0 | 5646 | -48.7 | -18.6 | 105.8 | -56.7 | 2 |
| | 3448 | -38.0 | -20.1 | 69.5 | 40.4 | -91.5 | 6406 | -65.2 | 9.5 | 0.7 | -55.6 | 3 |
| | 968 | -29.9 | -14.8 | 199.0 | 65.4 | -90.2 | 5015 | -40.2 | -21.2 | 58.8 | -55.5 | 4 |
| | 2710 | -35.3 | -21.9 | 99.8 | 42.3 | -89.6 | 6321 | -51.5 | 1.0 | -6.6 | -51.2 | 5 |
| | 5111 | -25.8 | -19.9 | 120.1 | 55.8 | -89.5 | 3448 | -38.0 | -20.1 | 69.5 | -51.1 | 6 |
| | 4119 | -52.0 | 10.5 | 60.1 | 53.8 | -89.3 | 4489 | -45.8 | -6.5 | 34.3 | -48.9 | 7 |
| | 3607 | -39.7 | -13.6 | 66.8 | 42.3 | -88.9 | 9940 | -56.3 | 3.1 | 49.3 | -48.3 | 8 |
| | 1238 | -35.8 | -12.2 | 95.9 | 48.1 | -86.4 | 2363 | -47.1 | -3.5 | 29.8 | -47.7 | 9 |
| | 9831 | -34.0 | 5.5 | 17.9 | 59.6 | -86.3 | 9056 | -24.8 | -27.0 | 44.3 | -47.3 | 10 |
| | 6321 | -51.5 | 1.0 | -6.6 | 34.6 | -85.8 | 2710 | -35.3 | -21.9 | 99.8 | -47.3 | 11 |
| | 3841 | -30.0 | -7.3 | 73.1 | 55.8 | -85.7 | 1285 | -42.3 | -7.6 | 28.7 | -47.1 | 12 |
| | 663 | -35.5 | -14.8 | 54.3 | 40.4 | -85.2 | 8005 | -39.2 | -10.3 | 23.6 | -47.1 | 13 |
| ***Chunk 4*** | **9109** | **-47.3** | **-24.4** | **120.3** | **87.5** | **-147.2** | 7608 | -44.7 | -38.9 | 74.3 | -76.2 | 1 |
| | 7608 | -44.7 | -38.9 | 74.3 | 62.5 | -138.7 | 1031 | -43.1 | -38.9 | 139.6 | -68.1 | 2 |
| | 4947 | -41.8 | -17.7 | 106.0 | 81.3 | -130.1 | 4971 | -45.3 | -24.8 | 40.8 | -66.0 | 3 |
| | 4971 | -45.3 | -24.8 | 40.8 | 62.5 | -128.5 | 1714 | -61.4 | -7.2 | 37.8 | -64.8 | 4 |
| | 1714 | -61.4 | -7.2 | 37.8 | 62.5 | -127.3 | 7602 | -46.6 | -20.3 | 45.5 | -62.4 | 5 |
| | 297 | -21.8 | -30.4 | 137.6 | 87.5 | -126.0 | 97 | -72.0 | 3.9 | 57.5 | -62.3 | 6 |
| | 9634 | -37.5 | -18.4 | 50.7 | 75.0 | -125.9 | 2971 | -63.1 | 0.8 | 17.2 | -60.6 | 7 |
| | 1177 | -30.1 | -21.9 | 74.9 | 81.3 | -125.8 | **9109** | **-47.3** | **-24.4** | **120.3** | **-59.7** | **8** |
| | 1031 | -43.1 | -38.9 | 139.6 | 56.3 | -124.3 | 1611 | -71.9 | 11.5 | 12.8 | -59.1 | 9 |
| | 600 | -41.8 | -26.7 | 159.7 | 68.8 | -121.3 | 857 | -64.2 | 2.4 | 31.3 | -58.7 | 10 |
| | 2964 | -44.8 | -17.5 | 102.1 | 68.8 | -120.9 | 2330 | -68.8 | 9.0 | 19.3 | -57.8 | 11 |
| | 8675 | -38.2 | -19.3 | 128.0 | 75.0 | -119.7 | 1706 | -63.3 | 4.2 | 13.2 | -57.8 | 12 |
| | 1341 | -32.6 | -18.2 | 77.8 | 75.0 | -118.0 | 6349 | -65.5 | 7.4 | 4.2 | -57.7 | 13 |

*Table S3: Selected ITPRIPL2 residues for docking constraints.*

| ***Residue*** | ***Amino acid*** |
|---|---|
| *12* | *F* |
| *15* | *L* |
| *19* | *L* |
| *23* | *L* |
| *26* | *L* |
| *27* | *Y* |
| *30* | *L* |
| *48* | *F* |
| *51* | *L* |
| *52* | *K* |
| *55* | *V* |
| *58* | *L* |
| *59* | *L* |
| *62* | *V* |
| *66* | *C* |
| *70* | *V* |

*Table S4: AlphaFill result scores including optimized scores for selected residues*

| ***Compound*** | ***PDBID*** | ***g-RMSd*** | ***Asym*** | ***l-RMSd*** | ***TCS*** | ***Optimized TCS*** |
|---|---|---|---|---|---|---|
| ***1SY (cGAMP)*** | *5VDP.A* | *2.93* | *G* | *1.85* | *0.49* | *0.03* |
| | *4LEZ.A* | *9.31* | *Q* | *1.95?* | *0.22* | |
| | *6MJX.A* | *13.00* | *K* | *0.51* | *0.71?* | *0.04* |
| | *4O67.A* | *13.12* | *D* | *2.34* | *0.73?* | |
| ***4BW*** | *5VDT.A* | *2.97* | *I* | *1.88?* | *0.58* | |
| ***5GP*** | *7BUJ.A* | *3.29* | *T* | *1.55?* | *0.34* | |
| ***9BG*** | *5VDQ.A* | *2.95* | *H* | *1.88?* | *0.65?* | |
| ***APC*** | *7UTT.A* | *3.34* | *V* | *1.04?* | *0.40* | |
| ***ATP*** | *6CTA.A* | *2.87* | *J* | *0.56* | *0.64* | *0.08* |
| | *4K97.A* | *11.13* | *N* | *0.71* | *0.72?* | |
| ***GTP*** | *7BUJ.A* | *3.29* | *S* | *3.02* | *0.74?* | |
| | *7BUJ.B* | *3.31* | *U* | *2.86* | *0.73?* | |
| | *7UXW.A* | *3.33* | *W* | *0.16* | *0.78?* | |
| | *7UYZ.A* | *3.34* | *Y* | *1.59?* | *0.15* | |
| | *7UYQ.A* | *3.34* | *X* | *1.04?* | *0.40* | |
| | *4K98.A* | *9.31* | *P* | *1.94?* | *0.31* | |
| ***ZN*** | *6O47.A* | *3.15* | *B* | *2.19* | *0.03* | |
| | *5XZE.A* | *5.45* | *R* | *1.52?* | *0.09* | |
| | *6NAO.A* | *6.09* | *L* | *3.98?* | *0.60* | |
| | *4KM5.A* | *8.93* | *C* | *4.82?* | *0.58* | |
| | *4K98.A* | *9.31* | *O* | *0.63* | *0.14* | |
| | *4O69.A* | *10.57* | *F* | *2.23* | *0.15* | |
| | *6NFG.A* | *12.42* | *M* | *5.92?* | *0.51* | |
| | *4O67.A* | *13.12* | *E* | *9.04?* | *0.27* | |
